# Antiviral T-cell Biofactory platform for SARS-CoV-2

**DOI:** 10.1101/2022.06.26.497669

**Authors:** Marvin A. Ssemadaali, Sherri Newmyer, Harikrishnan Radhakrishnan, Juan Arredondo, Harold S. Javitz, Satya Dandekar, Parijat Bhatnagar

## Abstract

Vaccines help reduce new infections, but interventions that can prevent the disease from transitioning to a severe stage are rather limited. Dysregulated IFN kinetics are mostly exploited by pathogenic viruses, including SARS-CoV-2. The clinical benefits of systemically infused IFN are, unfortunately, mired by undesired side effects. To address this situation, we engineered a T cell to synthesize interferons (IFNs) as antiviral proteins upon recognizing the virus envelop protein of SARS-CoV-2, i.e., *anti-SARS T-cell Biofactory*. The T-cell Biofactory, capable of regulating the IFN expression with spatiotemporal resolution within the infected tissues, can mitigate these concerns. In this work, we determined the prophylactic and therapeutic effects of the type-I and type-III IFNs produced from the T-cell Biofactory against SARS-CoV-2 infection in host cells and investigated the expression profiles of ensuing IFN-stimulated genes (ISGs). To enable the translation of T-cell Biofactory as an effective antiviral countermeasure, we also investigated an irradiation dose that renders the T-cell Biofactory non-proliferative and thus non-oncogenic. The ongoing public health crisis motivated us to direct the T-cell Biofactory technology to target SARS-CoV-2. The T-cell Biofactory, based on T cells engineered with chimeric antigen receptors (CAR T cells), is a platform technology that can be rapidly re-engineered and become available for targeting any new pathogen.

## 2.0 Introduction

Type-I and type-III interferon (IFN) response, mediated through pattern recognition receptors in host cells, is the first line of defense against viruses and acts by autocrine/paracrine signaling to induce hundreds of antiviral IFN-stimulated genes (ISGs)^1^. The dysregulated IFN response in SARS-CoV-2 infections indicates the progression of COVID-19 to different severity levels^2-8^. While both type-I and type-III IFNs generate similar ISG expression profiles, the timing and proportion of their expression differs during the infection and determines the severity of the disease. For example, the type-I IFN response is short-lived and occurs during early stages of the infection to serve as a prophylaxis^3,5^, and type-III IFNs prevent the progression of the disease to severe stages^9,10^ by exerting a long-lasting, non-inflammatory therapeutic response that helps clear systemic infection. Systemically delivered exogenous type-I IFN has shown therapeutic benefits when infused before the peak viral load; however, its use after the infection does not prevent the onset of severe stages^11-14^. Side effects like inflammation, tissue damage, and multiorgan failure have been observed when these IFNs are administered incorrectly^11,12,15^. Calibrated and timely delivery of exogenous IFN dosage, limited to the affected organs with spatiotemporal resolution, is a critical unmet need that would prevent the disease from transitioning to severe stages.

Previously we reported on engineering immune cells with chimeric antigen receptors (CAR) that, upon engaging antigen-presenting target cells, produce non-endogenous proteins for exerting the desired effect locally, i.e., T-cell Biofactory^16^ and NK-cell Biofactory^17^. Recently, we also directed this platform for antigen^18^ and serology testing^19^ for SARS-CoV-2. In the current study, we used this technology to develop two different classes of antiviral T-cell Biofactory that respectively synthesize type-I (IFN-α2^3^ or IFN-β1^5^) and type-III (IFN-λ2^10^ or IFN-λ1^9^) IFNs to compensate for the abnormal IFN biodistribution kinetics in viral infections. We determined the prophylactic and therapeutic effects of the IFNs produced by the antiviral T-cell Biofactory platform on SARS-CoV-2 infected cells and investigated the expression profiles of ISGs in the host cells in response to these IFNs. We also determined the radiation dose (γ-radiation) needed to render the T-cell Biofactory non-proliferative making the cellular products suitable for clinical use^17,20-22^, while conserving the cell-based IFN production. This process ensures safe administration of the T-cell Biofactory without potential oncogenesis.

## 3.0 Results

The artificial cell-signaling pathway, which comprises three constant (*Receptor, Actuator, Secretor*) and two variable (*Sensor, Effector*) domains arranged *in cis*^*16*^, was used to generate the T-cell Biofactory that, on detecting the Spike glycoprotein (Sgp) as an antigenic biomarker on the surface of host cells infected with SARS-associated coronaviruses (CoV), produces type-I or type-III IFNs with antiviral effects. The constant domains provide functionality to the T-cell Biofactory and include a transmembrane molecule [1: *Receptor*, part of the CAR] that mobilizes the T-cell activation machinery [2: *Actuator*] to express the desired transgene. To assist the secretion of IFNs, we used the natural signal peptide [3: *Secretor*] of the two IFNs and removed the one we previously standardized^16^. Variable domains can be exchanged to reprogram the T-cell Biofactory to specifically identify the diseases cells and exert the desired therapeutic effect to neutralize the pathology. For transforming the platform into the *anti-SARS T-cell Biofactory*, we tested several from among a range of antibodies that cross-react with SARS-CoV-2 Sgp, the results of which have been published elsewhere^18^. Our tests validated VHH-72 (PDB: 6WAQ) [1: *Sensor*, also a part of the CAR] as an effective candidate, a camelid-derived single-domain heavy chain that binds to the Sgp of SARS-CoV-1 and previously identified by others^23^ to cross-react with the Sgp of SARS-CoV-2 (**Figure S1**). The T-cell Biofactory platform can, however, be reprogramed for specificity toward a different viral pathogen by replacing the sequence for VHH-72 with a VHH or variable heavy-light [V_H_-V_L_] portion of the single-chain variable fragment [scFv] of different antibody, as we previously demonstrated by us^18^. On engaging the Sgp, the anti-SARS T-cell Biofactory mobilizes the T cell’s transcriptional machinery to synthesize type-I (IFN-α2, IFN-β1) or type-III (IFN-λ2, IFN-λ1) IFNs [2: *Effector*] reported to exert antiviral effects ^3,5,9,10.^ **Figure 1** shows the schematic and function of the anti-SARS T-cell Biofactory that identifies the SARS-CoV-2-specific Sgp on an infected host cell independent of its presentation in the peptide-major histocompatibility complex (pMHC).

**Figure 1.**
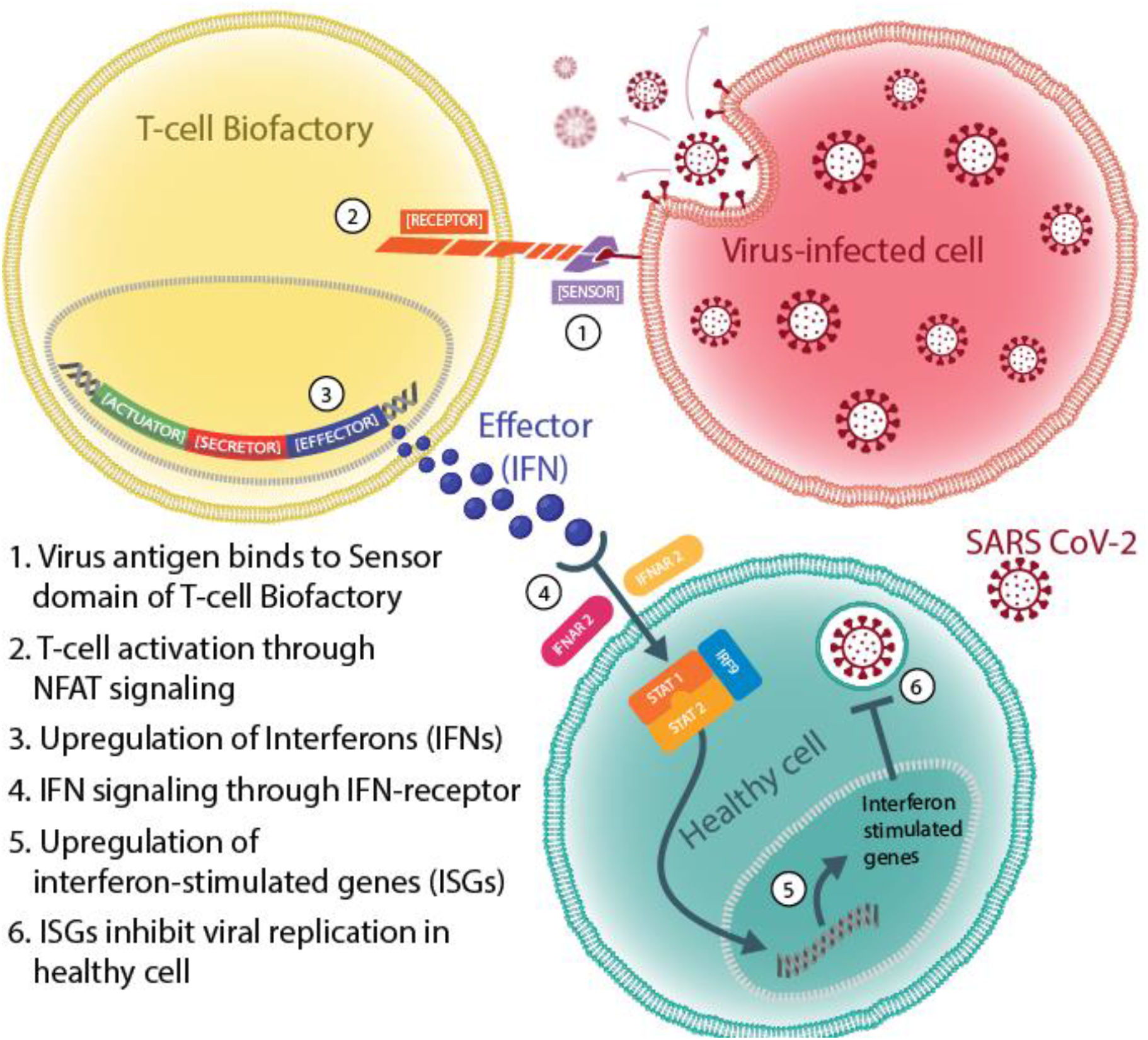
Schematic function of the anti-SARS T-cell Biofactory. The schematic illustrates the function and IFN production from anti-SARS T-cell Biofactory

The SARS-CoV-2-specific function of the anti-SARS T-cell Biofactory was assessed using two different types of Target-Cells. Initial tests were conducted using the HEK293T/17 cell line that is engineered to stably express the Sgp from SARS-CoV-2 as pseudo-infected Target-Cells (*SARS-CoV-2-Sgp-cell*) (**Figure 2A** and **2B**). We validated the results in a BSL3 containment facility by using infected Target-Cells (Vero-E6 host cells infected with competent SARS-CoV-2 virus) (**Figure 2C** and **2D**). Non-engineered HEK293T/17 or uninfected Vero-E6 parental cell lines, respectively, were used as the negative controls. We prepared two classes of the anti-SARS T-cell Biofactory that produced different Effector proteins: (i) type-I (IFN-α2; IFN-β1) and (ii) type-III (IFN-λ2; IFN-λ1). Our results guided the prioritization of IFN-β1 (type-I IFN candidate) and IFN-λ2 (type-III IFN candidate) for more detailed investigation. The results from IFN-α2 (type-I IFN) and IFN-λ1 (type-III IFN) are shown in supporting information (**Figure S2**). Our assessment determined that 20-Gy dose of the gamma radiation, which renders the T-cell Biofactory non-proliferative without degrading our artificial cell-signaling pathway, is non-oncogenic^17,22^ (**Figure S3**).

**Figure 2.**
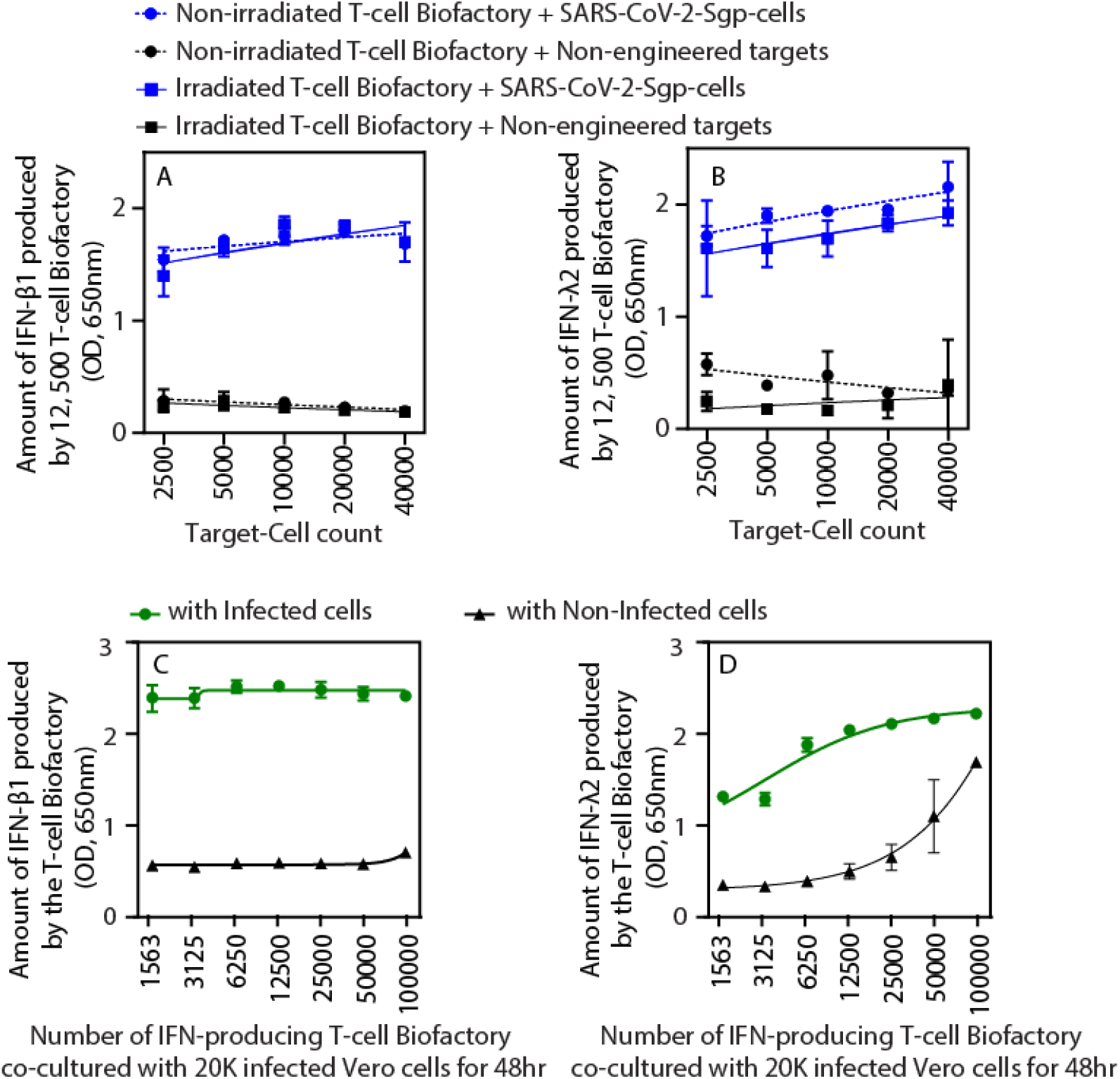
IFN production by the anti-SARS T-cell Biofactory (IFN-β1, IFN-λ2). T-cell Biofactory produce respective IFNs upon stimulation by the Target-Cells. Production of **(A)** IFN-β1 (type-I IFN) and **(B)** IFN-λ2 (type-III IFN) from the T-cell Biofactory (irradiated and non-irradiated) is proportional to the Target-Cell (SARS-CoV-2-Sgp-cells) count. 12,500 T-cell Biofactory were incubated with Target-Cells for 24 hours. Production of **(C)** IFN-β1 (type-I IFN) and **(D)** IFN-λ2 (type-III IFN) from the T-cell Biofactory is proportional to the number of T-cell Biofactories when stimulated by SARS-CoV-2-infected Vero-E6 Target-Cells. 20,000 Vero-E6 cells, infected with SARS-CoV-2 at MOI of 0.5, were used to stimulate the T-cell Biofactory for 48 hours. For all observations, n = 3, error bars indicate ±1 standard deviation (SD).

**Figures 2A** and **2B** demonstrate the expression of type-I IFNs (IFN-β1) and type-III IFNs (IFN-λ2) by the T-cell Biofactory respectively and compares the results with the IFNs produced from the two T-cell Biofactories irradiated at 20 Gy. Consistent with our previous work on the anti-tumor T-cell Biofactory^16^ and the anti-tumor NK-cell Biofactory^17^, the Effector protein (IFNs) expression was proportionate to the Target-Cell count and was observed in all Effector-Cell to Target-Cell (E:T) ratios. The IFN expression by the T-cell Biofactory was significantly elevated (p<0.00005 at all E:T) when stimulated by the target SARS-CoV-2-Sgp-cells, compared to when it was stimulated by the non-engineered negative control cells. Similar results were observed when both T-cell Biofactories were irradiated with 20 Gy (p<0.01 at all E:T).

To further validate our observations made in the T-cell Biofactory with pseudo-infected target SARS-CoV-2-Sgp-cells, we exchanged the engineered target SARS-CoV-2-Sgp-cells with Vero-E6 host cells infected with competent SARS-CoV-2 virus (Isolate: Hong Kong/VM20001061/2020) in a BSL3 containment facility. The results presented in **Figures 2C** and **2D** demonstrate that type-I IFN (IFN-β1) and type-III IFN (IFN-λ2) expression by the respective anti-SARS T-cell Biofactories was significantly higher compared to the IFN expression from the T-cell Biofactories stimulated by non-infected Vero-E6 cells (p<0.00005 for IFN-β1 and p<0.01 for IFN-λ2 at all E:T ratios). Evidence of the broad applicability of anti-SARS T-cell Biofactory is once again shown in **Figure S4**. This is attributed to the use of VHH-72 as an antigen-binding domain (Sensor) that has specificity towards SARS-CoV-1 Sgp^23^. As shown in **Figure S4**, IFN production from the anti-SARS T-cell Biofactory was observed when stimulated by the pseudo-infected Target-Cells presenting SARS-CoV-1-specific Sgp (*SARS-CoV-1-Sgp-cell*).

Figure 3. reports on the protective effects of the type-I IFNs (IFN-β1) and type-III IFNs (IFN-λ2) secreted by the T-cell Biofactory. To simulate the role of timing in infection^24^, we investigated the relative protection offered by the two IFNs before and after the viral challenge, i.e., their prophylactic and therapeutic effects. Toward this goal, we either pretreated Vero-E6-Luc2^+^ host cells with the IFN-containing supernatants from the respective activated T-cell Biofactories (type-I IFN-β1 or type-III IFN-λ2) before infecting them with SARS-CoV-2 (prophylactic effect), or we co-cultured the two T-cell Biofactories with previously infected Vero-E6-Luc2^+^ host cells (therapeutic effect). We assessed the viability of host cells to determine the protection achieved in both cases. **Figure S5** shows the protective effects from the other T-cell Biofactories (type-I IFN-α2 and type-III IFN-λ1).

**Figure 3.**
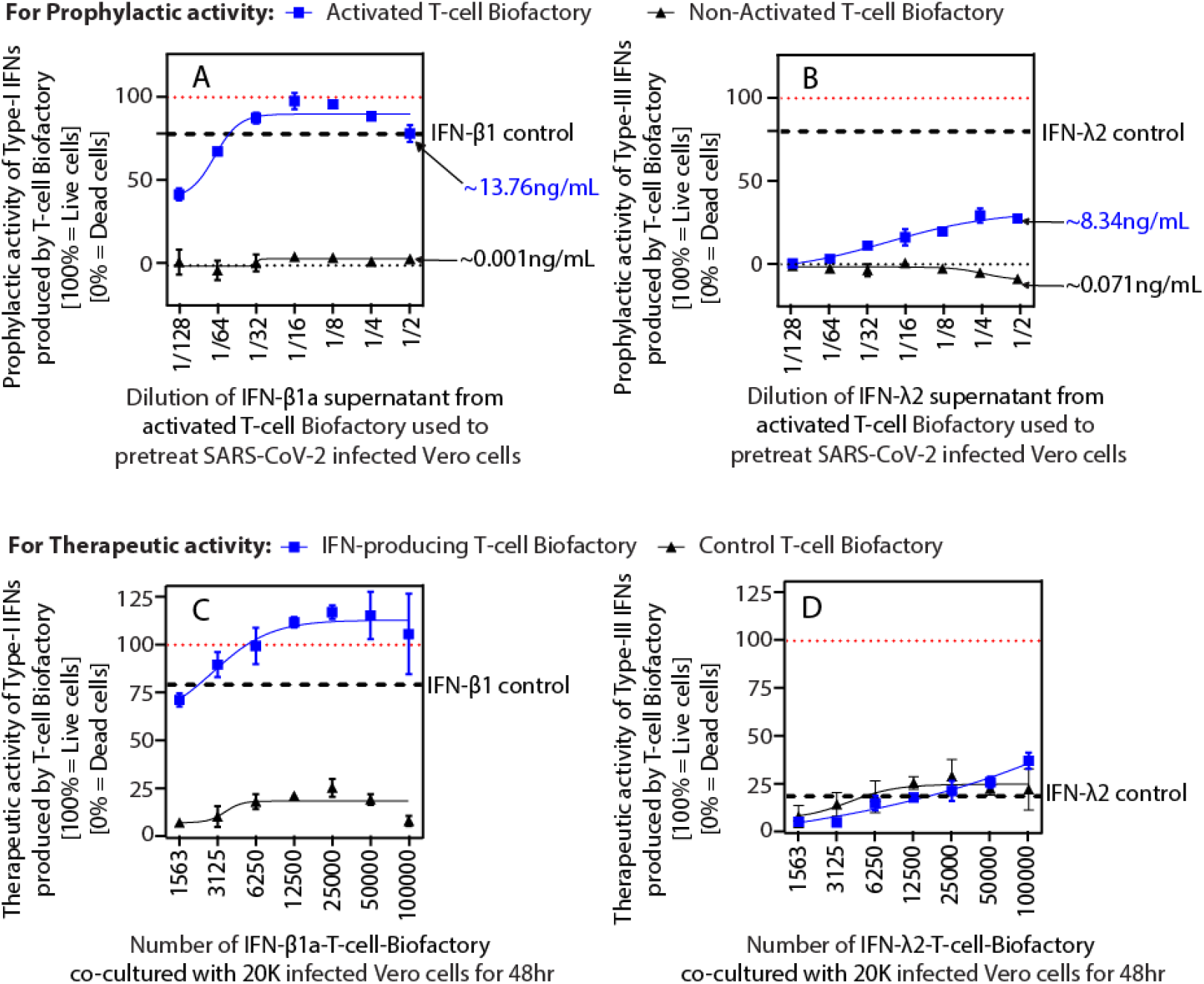
Prophylactic and therapeutic activity of the anti-SARS T-cell Biofactory (IFN-β1, IFN-λ2). The IFN produced by the T-cell Biofactory can be used as a prophylaxis **(A, B)** and a therapeutic **(C, D)** against SARS-CoV-2 infection. The protective function of the T-cell Biofactory is proportional to the amount of IFN produced **(A, C)** IFN-β1 (type-I IFN) and **(B, D)** IFN-λ2 (type-III IFN). For **(A, B)** MOI=0.1; **(C, D)** MOI=0.5. All data were collected with 20,000 Vero-E6 cells incubated with T-cell Biofactories for 48 hours. Respective recombinant human IFN at 1 μg/mL were used as controls. For all observations, n = 3, error bars indicate ±1 SD.

Results in **Figure 3A** and **3B** illustrate the prophylactic activity of type-I IFNs (IFN-β1) and type-III IFNs (IFN-λ2) produced by the respective activated T-cell Biofactories when compared to the negative control from non-activated T-cell Biofactories. Statistically significant protection (p<0.001) was observed at all IFN-β1 concentrations (**Figure 3A**). While IFN-λ2 was still protective at all IFN dilutions less than 1/32 (i.e., concentrations above 0.587 ng/mL) (p<0.005) (**Figure 3B)**, this ability was lower compared to that of IFN-β1. Results presented in **Figure 3C** and **3D** demonstrate the therapeutic activity of type-I IFN (IFN-β1) and type-III IFN (IFN-λ2) T-cell Biofactories when co-cultured with infected host cells. A T-cell Biofactory without Actuator/Secretor/Effector domains was used as a negative control. As expected, the therapeutic protection offered by the IFN-producing T-cell Biofactories was proportional to their number (indicating the amount of IFN produced^16,17^) (see **Figure 3C** for IFN-β1 and **Figure 3D** for IFN-λ2). The therapeutic effect demonstrated by the type-I IFN-β1-producing T-cell Biofactory was higher (p<0.005 at all Effector-Cell counts) when compared to the control T-cell Biofactory (**Figure 3C**). Although the effect was not significant with the type-III IFN-λ2-producing T-cell Biofactory, an effect trend was observed (**Figure 3D**). This prophylactic and therapeutic activity demonstrates the potency of the IFNs produced by the T-cell Biofactory and confirm reports from previous studies that the SARS-CoV-2 virus is susceptible to IFN treatment^4,25^.

To validate the antiviral effect of IFN-β1 produced by the T-cell Biofactory in **Figure 3**, we analyzed the regulation of genes in the type-I IFN signaling pathway in Vero-E6 host cells using the NanoString Host Response Panel analysis and benchmarked it against treatment with commercially available IFN (**Figure S6**). The results were validated using quantitative PCR by assessing the expression profiles of a few representative ISGs (ISG15, Mx1, OAS1, IFIT1, IFI44, RSAD2, RNASEL), shown in **Figure 4**. Over-expression of these ISGs in host cells inhibits viral entry, replication, and viral budding or egress^1,26,27^, and, therefore, prevents viral spread. **Figure 4** also includes data on the STAT1 gene indicating the upregulation of the Janus Kinase-Signal Transducers and Activators of Transcription (JAK-STAT) signaling pathway^1^, activated by both type-I and type-III IFNs, that regulates the expression of ISGs for controlling viral replication. The results in **Figure 4A – 4H** regarding type-I IFN signaling in Vero-E6 and its effect on the downstream expression of ISGs also substantiates the antiviral effects observed in **Figure 3**. Similar expression profiles were observed when the supernatants from the IFN-β1-producing T-cell Biofactory were used to treat Calu-3 human epithelial cells (**Figure S7**).

**Figure 4.**
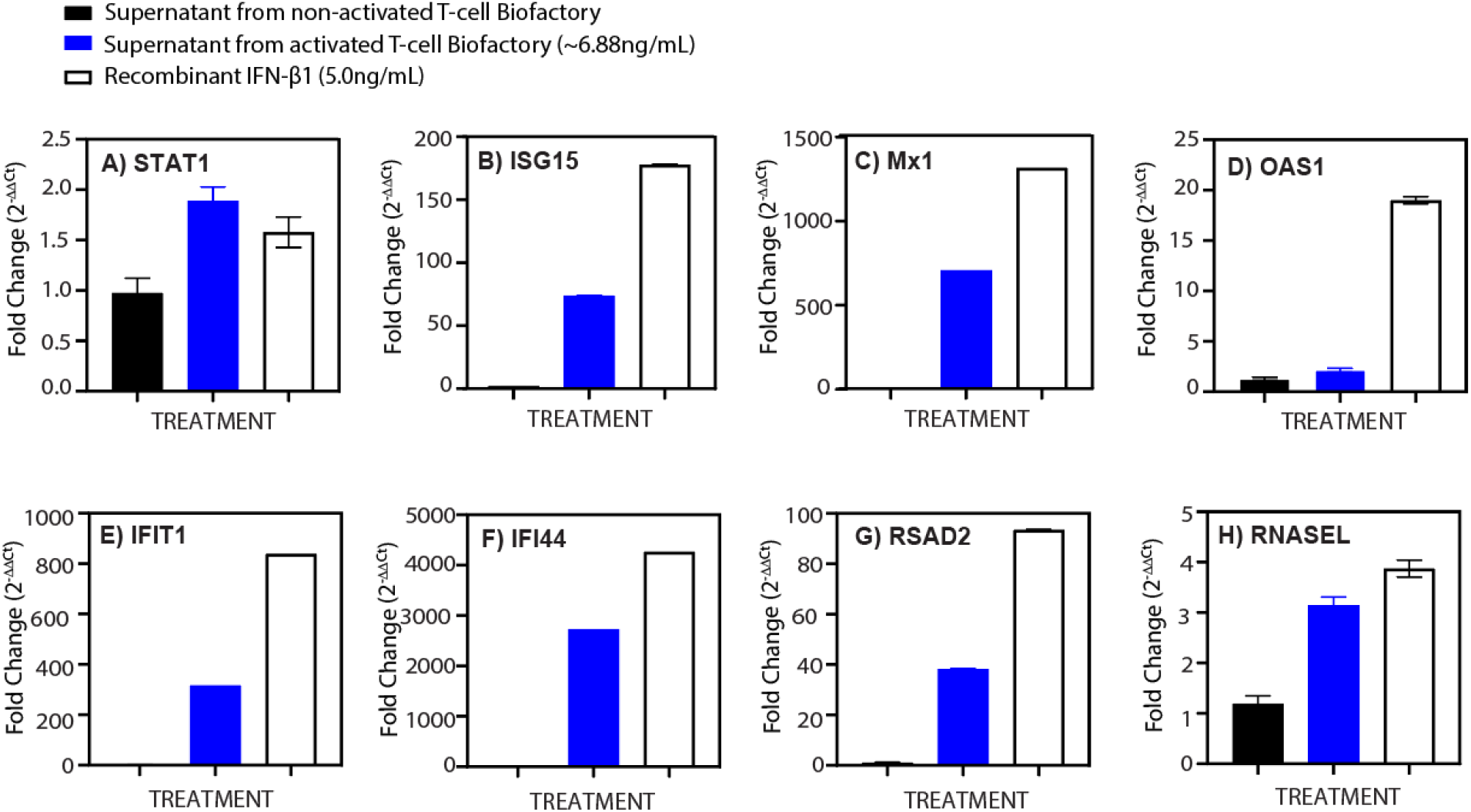
Effect of IFN produced by the anti-SARS T-cell Biofactory on Vero-E6 host cells. IFN-β1 (type-I IFN) produced by the T-cell Biofactory upregulates the IFN signaling pathway, and its effect is similar to the exogenously delivered recombinant human IFN-β1 (control). Vero-E6 cells were treated with the supernatant produced by the IFN-β1-producing T-cell Biofactory or control for 24 hours. For all observations, n = 3, error bars indicate ±1 standard errors of the mean (SEM).

## 4.0 Discussion

Vaccines prime adaptive immunity by inducing a convalescent stage and are undoubtedly effective as a prophylaxis. Dearth of antiviral treatments to stop the transition of disease to severe stages, however, present an important gap in the currently available clinical options. The use of antiviral drugs^28,29^ and antibody-based therapies^30,31^ reduce instances of severe disease, but the threat of drug resistance among new variants that continue to emerge is driving the search for better alternatives. The IFN signaling system, which exerts antiviral effects through the induction of hundreds of ISGs, is an effective option that offers to clear most viruses at the incubation and prodromal stages at all degrees of severity. Using IFNs to treat viral infections could also circumvent recent concerns reported with some antibody therapies and drugs^32,33.^ IFN therapy, however, remains untapped because the pathogens evolve in vivo and disease progresses during the infection, making it hard to determine the effective but still safe IFN dosage. The difficulty of correcting for the misalignment between the bioavailability of systemically delivered IFN and the localized disease microenvironment has been responsible for adverse effects such as inflammation, tissue damage, and multiorgan failure^11,12,15^. Our results illustrate (a) the calibrated production of two different IFNs (type-I IFN-β1 and type-III IFN-λ2) from two different T-cell Biofactories, (b) the protection they provide to SARS-CoV-2 infected host cells, and (c) the upregulation of antiviral ISGs in response to these IFNs.

Recent outbreaks have flagged a known fact that such events have been common throughout history and are likely to happen again^42,43^. Like most other pathogenic viruses^34,35^, SARS-CoV-2 evades the host’s IFN immune response, which is critical in the establishment of an early antiviral defense^15,36^. It has been reported that SARS-CoV-2 causes a delayed IFN response that contributes to the severity of COVID-19^13,14.^ Although clinical application in COVID-19 patients is still being evaluated, several studies have demonstrated that recombinant IFNs can reduce viral replication in vitro^6,25^, and in preclinical animal models^9,37.^ Clinical trials for both type-I and type-III IFNs have also been undertaken in COVID-19 patients to demonstrate safety and efficacy^38-41^. However, these did not meet the required success criteria due to lack of control on the IFN dosage and its timing with respect to the disease stage^24,26,40,42.^ The T-cell Biofactory has been designed to mitigate these exact concerns by presenting a unique approach to administering IFNs. It uses cues from the infected target cells to regulate its transcriptional machinery, resulting in the synthesis of calibrated IFN amounts with spatiotemporal resolution. The innovation in antiviral T-cell Biofactory platform is that it employs an alternative synthetic pathway that bypasses the natural pathway of triggering the IFN signaling through pattern-recognition receptors, and is often compromised in pathogenic viral infections^26,34,35^. This approach is expected to reduce inefficacies due to suboptimal IFN levels and hyperinflammation caused by excess infusion, both of which have been reported as adverse effects of systemic infusions^23,25,39,41^. The anti-SARS T-cell Biofactory adds to the growing portfolio of cell-based interventions for viral diseases^43-45^.

Nevertheless, challenges need to be overcome before the antiviral T-cell Biofactory can advance toward the clinical use. Some of these include: (i) developing non-exhausting cells for long-term systemic circulation, (ii) assessing their extravasation to the infection sites, (iii) exploring a universal cell chassis for allogeneic transplant, and (iv) conducting preclinical *in vivo* trials to validate these effects. The benefits of this work will also extend beyond the SARS-CoV-2. This is because an antibody with pan-CoV specificity^46^, many of which are currently in development^47^, offers an opportunity to develop a pan-CoV T-cell Biofactory that, in the event of another CoV pandemic. The platform can also be genetically reprogrammed to target other pathogenic viruses, as previously described by us^18^. In addition, the inclusion of allogeneic options^48-51^ among the cell chassis will further the impact of this work. Our approach therefore represents a substantive departure from the status quo and offers a powerful means for curtailing future pandemics.

## 5.0 Acknowledgements

The research work reported in this publication was supported in part by the National Institute of Biomedical Imaging and Bioengineering (DP2EB024245: NIH Director’s New Innovator Award Program (https://commonfund.nih.gov/newinnovator); the National Cancer Institute (R21CA236640, R33CA247739) of the National Institutes of Health (NIH); and the Defense Advanced Research Projects Agency (DARPA) (D19AP00024: DARPA Young Faculty Award (https://www.darpa.mil/work-with-us/for-universities/young-faculty-award). The content is solely the responsibility of the authors and does not necessarily represent the official views of the NIH or DARPA. The authors thank Didier Trono (Ecole Polytechnique Fédérale de Lausanne, Lausanne, Switzerland) for providing the lentivirus packaging plasmids; Mary Lanier (SRI International) for providing the Vero-E6 cells and SARS-CoV-2 virus culture; Nancy Craig (Johns Hopkins University School of Medicine) for providing the piggyBac Transposase sequence; Praveer Sharma (NanoString) for his help with the NanoString nSolver analysis; and Julie Thomas (SRI International) for manuscript proofreading and editing.

## 6.0 Author contributions

PB, the principal investigator, conceptualized the anti-SARS T-cell Biofactory, designed the experimental investigations, and wrote the manuscript. MS designed the experimental investigations, performed the assays, and wrote the manuscript. HSJ was the co-investigator and designed the statistical tests. SD advised on experimental designs, and JA, SN, and HR conducted experiments. All authors read and contributed to preparation of the manuscript.

## 8.0 Experimental section

### Materials and reagents

Engineered Jurkat E6-1 (ATCC, Cat# TIB-152) cell lines were maintained in complete RPMI media (RPMI1640 [Corning, Cat #10-040-CV], 10% were heat-inactivated fetal bovine serum or FBS [Sigma-Aldrich, Cat #F2442-500ML], and 1X Penicillin-Streptomycin solution [Corning, Cat #30-002-Cl]). Parental and engineered Vero-E6 cells (ATCC, Cat # CRL-1586) were cultured in complete EMEM (EMEM growth media [Corning, Cat #10-009-CV] supplemented with 10% heat-inactivated FBS and 1X Penicillin-Streptomycin solution). Parental and Engineered HEK293T/17 cells (ATCC, Cat #CRL-11268) were cultured in complete DMEM (DMEM growth media [Corning, Cat #10-013-CV] supplemented with 10% FBS and 1X Penicillin-Streptomycin solution). All cells were expanded, and liquid nitrogen stocks were maintained using freezing media (50% FBS, 40% growth media and 10% Dimethyl sulfoxide). Plasmids encoding different genetic payloads (transfer plasmids) were designed in SnapGene software (GSL Biotech LLC) and sub-cloned into lentivirus vector plasmid (System Biosciences, Cat #CD510B-1) or PiggyBac Transposon vector plasmid (System Biosciences, Cat # PB510B-1). Plasmids encoding 2^nd^-generation packaging plasmids (psPAX2 – Cat #12260, pMD2.G – Cat #12259) were obtained from Addgene. pAdvantage was obtained from Promega (Cat #E1711). PiggyBac Transposase (i7pB) sequence was provided by Dr. Nancy Craig at the Johns Hopkins University School of Medicine ^52^. An insert for the EF1alpha promoter – i7pB transgene –bGH poly(A) signal was chemically synthesized and assembled using overlapping PCR products into pUC19 (GenBank: L09137, New England Biolabs, #N3041). All plasmid preparation services (chemical synthesis of DNA insert sequences, sub-cloning into respective vector backbones, and the amplification) were obtained from Epoch Life Science, Inc. (Missouri City, TX). For lentivirus production, Transporter 5™ reagent (Polysciences, Inc, Cat #26008-5) was used to transfect parental HEK293T/17 cells. The collected lentivirus was transduced into Jurkat cells using Polybrene (abm®, Cat #G062). TransIT®-2020 transfection reagent (Mirus #MIR5400) was used to transfect PiggyBac Transposon system plasmids into parental HEK293T/17 cells to engineer stable antigen-presenting cells (APCs) or pseudo-infected host target cells. Puromycin dihydrochloride (ThermoFisher Scientific, Cat# A1113803) was used to select stable cells. Phosphate-buffered saline (PBS) without Ca^+2^ and Mg^+2^ (Corning, Cat #21-040-CV) was used to wash cells. Dr. Mary Lanier at SRI International provided the SARS-CoV-2 virus culture (BEI Resources, NIH; Hong Kong/VM20001061/2020 [Cat# NR-52282]). The viability of Vero-E6-Luc2^+^ cells was assessed using either CellTiter-Glo™ Luminescent Cell Viability Assay Kit (Promega, Cat# PR-G7570) or One-Glo® assay (Promega, Cat #E6110). HEK-Blue IFN-α/β cells (InvivoGen, Cat# hkb-ifnab) or HEK-Blue IFN-λ cells (InvivoGen, Cat# hkb-ifnl) were used to quantify the amount of IFNs produced by the T-cell Biofactory. Recombinant human interferon (rhIFN) proteins from R&D systems (IFN-β [Cat# 8499-IF-010/CF], IFN-λ1 [Cat#1598-IL-025/CF], IFN-λ2 [Cat#8417-IL-025/CF]) and IFN-α2 from InvivoGen [Cat# rcyc-hifna2b] were used as controls. RNA preparations used Direct-zol RNA Mini-Prep Kit (Zymo Research, Cat# 11-331) and reverse transcription by SuperScript III RT (Invitrogen, Cat# 18080044). TaqMan Universal PCR Master Mix (Applied Biosystems, Cat# 4305719) was used for gene expression qPCR assays.

### Generation of the anti-SARS T-cell Biofactory (Effector-Cell, E)

Lentivirus particles were prepared by packaging the transfer plasmid using 2^nd^ generation lentivirus system as detailed previously^53^. The Jurkat E6-1 suspension cell line was engineered with lentivirus particles carrying the genetic construct for the T-cell Biofactory. The V_H_-V_L_ sequence for FRa or MSLN binding *Sensor* domain, as used previously^16,17^, was exchanged with the sequence for VHH-72 (PDB: 6WAQ) and Nluc sequence was exchanged with the respective IFN as the *Effector* domain. Briefly, the cells were transduced with transgenes using lentivirus in the presence of 8 μg/mL Polybrene. After 48 hours, the engineered cells were placed under selection using 0.5 μg/mL of Puromycin dihydrochloride. The unmodified parental cell line was also placed under selection as a positive control for cell killing by Puromycin dihydrochloride. Following selection, cells were expanded as required for different assays and frozen using freezing media. For irradiation, the T-cell Biofactory was exposed to 20 Gy (or as indicated) using a ^137^Cs γ-emitting irradiator, Mark I-68A (JL Shepherd and Associates) at a dose rate of 222 mGy/min. The control cells were treated similarly except for the irradiation.

### Generation of SARS-CoV-2-Sgp-cell and Vero-E6-Luc2^+^ (Target-Cells, T)

Parental HEK293T/17 or Vero-E6 cells were engineered with plasmids using PiggyBac transposon vector backbone, as previously reported ^54^, to express Sgp from SARS-CoV-2 (GenBank: QHD43416.1) or Luc2 Reporter transgene (GenBank: AY738222.1), respectively. A monolayer of cells (HEK293T/17 or Vero-E6) was transfected with the transposon plasmid (carrying the gene of interest) and transposase plasmid, in a ratio of 2.5:1, respectively, using TransIT®-2020 transfection reagent. After 48 hours of transfection, the transfected cells were placed under selection using Puromycin dihydrochloride (0.5 μg/mL for HEK293T/17; 3 μg/mL for Vero-E6-Luc2^+^). The unmodified parental cell lines were placed under selection as a positive control for cell killing by Puromycin dihydrochloride. The generated stable cell lines were expanded as required for different assays.

### Production of the IFNs (Effector) from the anti-SARS T-cell Biofactory

Effector-Cells (E) (irradiated or non-irradiated T-cell Biofactory) and Target-Cells (T) (SARS-CoV-2-Sgp-cell or non-engineered cells) were co-cultured at different E:T ratios in 100 μL of complete RPMI media in a single well of a 96-well plate. After the specified amount of time in co-culture, the amounts of IFNs produced by the T-cell Biofactory were assessed using the HEK-Blue IFN-α/β or IFN-λ reporter assays (see *IFN quantification assays*).

### Prophylactic effect of the IFNs produced by the anti-SARS T-cell Biofactory

2×10^5^ Target-Cells (SARS-CoV-2-Sgp-cells or non-engineered cells) were co-cultured with 1×10^6^ Effector-Cells (anti-SARS T-cell Biofactory) (E:T = 5:1) in 0.5 mL of complete RPMI media in a single well of a 24-well plate. After the specified amount of time in co-culture, the IFN-containing supernatants were collected and serially diluted (2-fold) in complete EMEM media. 100 μL of serially diluted IFN-containing supernatants was then used to pre-treat a monolayer of Target-Cells (Vero-E6-Luc2^+^ cells) for 24 hours (20,000 cells/well, triplicates) in a 96-well plate; recombinant human IFNs (1 μg/mL) were included as controls. The pre-treated cells were then infected with SARS-CoV-2 virus culture at a multiplicity of infection (MOI) of 0.1 for 48 hours. Cell viability in Vero-E6-Luc2^+^ cells was determined using the CellTiter-Glo™ Luminescent Cell Viability Assay Kit (see *Luminescence assays*). Live-dead status of the cells was indicated by normalizing the luminescence signal that indicates the ATP levels in live cells (100% = no virus infection, 0% = virus infection). The original amount of IFNs produced by each T-cell Biofactory was assessed using the HEK-Blue IFN-α/β or IFN-λ reporter assays (see *IFN quantification assays*).

### Therapeutic effect of the IFNs produced by the anti-SARS T-cell Biofactory

A monolayer of Vero-E6-Luc2^+^ (20,000/well in a 96-well plate) was infected with SARS-CoV-2 virus culture at an MOI of 0.5 and incubated for 2 hours to allow virus attachment. After 2 hours, the virus inoculum was removed, and 150 μL of serially diluted IFN-producing T-cell Biofactory (2-fold) was immediately added to the wells (triplicates). Recombinant human IFNs (1 μg/mL) and a T-cell Biofactory with Sgp-specific Sensor domain but no Actuator/Secretor/Effector domains (or non-infected Vero-E6-Luc2^+^ cells) were used as controls. After 48 hours of co-culture, 50 μL of IFN-containing supernatant was removed from each well to quantify the amount of IFNs produced by the T-cell Biofactory at each E:T ratio, using the HEK-Blue IFN-α/β or IFN-λ reporter assays (see *IFN quantification assays*). Then, Vero-E6-Luc2^+^ cell viability was determined by assessing Luc2 activity using the One-Glo® assay kit (see *Luminescence assays*). Live-dead status of the cells was indicated by normalizing the luminescence signal that indicates the ATP levels in live cells (100% = no virus infection, 0% = virus infection).

### Luminescence (Nluc®, CellTiter-Glo®, One-Glo®) assays

Luminescence assays were conducted by following the manufacturer’s protocol. Briefly, the substrate for enzymes (Nluc, CellTiter Glo, One-Glo) was diluted in the cell lysis buffer provided with the respective assay kit and added to the co-cultures in a 96-well plate. Following a brief incubation period (3 min for Nluc or 10 min for CellTiter Glo and One-Glo), bioluminescence was read on a microplate reader (Perkin Elmer, EnVision™ Multilabel Plate Reader Model: 2104-0010A).

### IFN quantification assays

The IFN production was assessed using HEK-Blue IFN-α/β and HEK-Blue IFN-λ reporter cells, following the manufacturer’s instructions. Briefly, in a single well of a 96-well plate, 150 μl of complete DMEM containing 50,000 HEK-Blue IFN-α/β and HEK-Blue IFN-λ reporter cells were mixed with 50 μL of IFN-containing supernatant from the activated T-cell Biofactory. Serial dilutions (10-fold) of recombinant human type I or type III IFNs (rhIFNs) in complete DMEM were added in parallel to generate a standard curve. After 24 hours of incubation, 20 μL of supernatants from the reporter cells (HEK-Blue IFN-α/β or HEK-Blue IFN-λ) were added to 180 μL of Quanti-blue substrate (InvivoGen) and incubated at 37°C for 1-2 hours. Absorbance was measured at 650 nm using a microplate reader (Perkin Elmer, EnVision™ Multilabel Plate Reader). The standard curves were then used to estimate the IFN concentrations produced by the respective anti-SARS T-cell Biofactory.

### Quantification of IFN-stimulated gene (ISG) mRNA expression

IFN-β1 supernatants from the stimulated T-cell Biofactory was diluted in complete EMEM media at 1:4 dilution ratio (∽6.88 ng/mL). In a 24-well plate, the diluted IFN supernatant was used to treat a monolayer of 2×10^5^ Vero-E6 (or Calu-3) cells (in triplicate) for 24 hours (5 ng/mL of recombinant IFN-β1 was used as a control). 300 μL of TRIzol was used to lyse cells, and RNA purifications were performed using a Direct-zol RNA Miniprep kit following manufacturer’s instructions. Purified RNA was reverse transcribed into cDNA using the SuperScript™ III Reverse Transcriptase. The cDNAs were analyzed by qPCR using TaqMan Universal PCR Master Mix and TaqMan gene expression assays. The following TaqMan primer/probe sets (obtained from Thermo Fisher Scientific) were used to assess type-I IFN signaling: GAPDH (Cat# Hs02786624_g1), 18S (Cat# Hs99999901_s1), ACTB (Cat# Hs03023880_g1) IFIT1 (Cat# Hs03027069_s1), IFI44 (Cat# Hs00951348_m1), STAT1 (Cat# Hs00234829_m1), ISG15 (Cat# Hs01921425_s1), OAS1 (Cat# Hs05048921_s1), RNASEL (Cat# Hs05030865_s1), RSAD2 (Cat# Hs04967697_s1), and MX1 (Cat# Hs00895608_m1). All qPCR was performed in 384-well plates and run on a ViiA7 real time PCR system (Cat# 4453545). ISG expression was calculated using the ΔΔCT method ^55^ by normalizing the threshold cycle (Ct) values to reference genes (GAPDH, 18S, and ACTB), and expressions are represented as fold changes over untreated cell samples.

### NanoString gene expression methods and data analysis

IFN-β1 supernatants from the stimulated T-cell Biofactory was diluted in complete EMEM media at 1:4 dilution ratio (∽6.88 ng/mL). In a 96-well plate, the diluted IFN supernatant was used to treat a monolayer of 1×10^5^ Vero-E6 cells (in triplicate) for 24 hours (5 ng/mL of recombinant IFN-β1 was used as a control). Cells were then collected in 20 μL of RLT Buffer (Cat# 79216, Qiagen Inc). Service for quantification of gene mRNA and differential expression analysis was obtained from NanoString Technologies (Seattle, WA, USA) using NanoString nCounter Host Response Panel. Detailed information and the gene list are available on NanoString official website at https://nanostring.com/products/ncounter-assays-panels/immunology/host-response/. The nSolver™ 4.0 analysis software (NanoString Technologies) was used to process the raw data that included quality control of the data and its normalization with respect to 5 house-keeping reference genes. Advanced Analysis Module in the nSolver™ analysis software and the Limma package in the R Statistical Computing Environment was used for data analysis and generating differential gene expression plots, pathway scores plots and heatmap. Differential gene expression between the treatment groups was determined using a variance-stabilized t-test in nSolver™.

### Statistical analysis

GraphPad Prism 9.2.0 (GraphPad Software, Inc) (for all figures except Figure S6) and nSolver™ analysis software (NanoString Technologies) (for Figure S6) was used to conduct all statistical analyses. The experimental design and logistical models used for each panel in the figures is described further in the Supporting Information.

