## Supplementary Information for "Antiviral T-cell Biofactory platform for SARS-CoV-2"

**Running title**: *Anti-SARS T-cell Biofactory*

**Supplemental statistical analysis**: For certain graphs, a four-parameter logistic model was used, i.e., $Y = Y_{min} + \{Y_{max} – Y_{min}\} / \{1 + 2^\wedge[b^* (\log_2[X_{50}] – X)]\}$, where Y is the value of ordinate at X; $Y_{max}$ and $Y_{min}$ are estimated parameters defining upper and lower asymptotes, respectively; b is a "Hill" parameter defining the slope at the inflection point of the fitted curve; and $X_{50}$ is an estimated parameter representing the X value corresponding to $(Y_{max} - Y_{min})/2$. For all line graphs, comparison of all data points was calculated using the false discovery rate (FDR) two-stage step-up multiple comparison procedure of Benjamini, Krieger, and Yekutieli with FDR = 1%. Error bars in these graphs extend 1 SD above and below the mean.

(i) *Figures 2A and 2B (IFN production by the irradiated and non-irradiated T-cell Biofactory when activated by pseudo-infected target cells).* The IFN production in the anti-SARS T-cell Biofactory stimulated by the target (SARS-CoV-2-Sgp-cell) or non-target (non-engineered) cells was fitted using the linear model Y = a + b * X where Y is the IFN amount produced, X is the $\log_2$ of the target cell count, a is the Y-intercept, and b is the slope of the fitted curve.

(ii) *Figures 2C and 2D (IFN production by the T-cell Biofactory when activated by virally infected target cells).* The IFN production in the anti-SARS T-cell Biofactory stimulated by the infected target or non-infected target cells, was fitted using a four-parameter logistic model, where the independent variable X is the $\log_2$ of the Effector-Cell count.

(iii) *Figures 3A and 3B (Prophylactic activity of IFNs produced by the T-cell Biofactory).* The ATP activity was normalized and fitted using a four-parameter logistic model, where the independent variable X is the $\log_2$ of the IFN-dilution.

(iv) *Figures 3C and 3D (Therapeutic activity of IFNs produced by the T-cell Biofactory).* The Luc2 activity, when T-cell Biofactory was co-cultured with infected or non-infected Vero-E6-Luc2+ target cells, was fitted using a four-parameter logistic model, where the independent variable X is the $\log_2$ of the Effector-Cell count.

(v) *Figures 4A – 4H (IFN signaling in Vero-E6 cells treated with type-I IFNs from activated T-cell Biofactory).* Differential gene expression was calculated using the ΔΔCT method by normalizing the sample Ct values to the mean Ct values of 3 reference genes, and expressions are represented as fold changes over untreated cell samples. Error bars represent standard errors of the mean (SEMs) from the three biological replicates.

(vi) *Figure S1 (Anti-SARS T-cell Biofactory specificity to SARS-CoV-2 and SARS-CoV-1 infections).* The Nluc reporter activity in the T-cell Biofactory stimulated by target cells (SARS-CoV-2-Sgp-cell and SARS-CoV-1-Sgp-cell) or non-target (non-engineered) cells was fitted using a four-parameter logistic model, where the independent variable X is the $\log_2$ of the Target-Cell count.

(vii) *Figures S2A and S2B (IFN production by the irradiated and non-irradiated anti-SARS T-cell Biofactory when activated by pseudo-infected target cells).* The IFN production in the T-cell Biofactory stimulated by the target (SARS-CoV-2-Sgp-cell) or non-target (non-engineered) cells was fitted using the equation $Y = a + b * X$ where Y is the IFN amount produced, X is the $\log_2$ of the Target-Cell count, a is the Y-intercept, and b is the slope of the fitted curve.

(viii) *Figures S2C and S2D (IFN production by the anti-SARS T-cell Biofactory when activated by* *virally infected target cells).* The IFN production in the T-cell Biofactory stimulated by the infected target or non-infected target cells was fitted using the using a four-parameter logistic model, where the independent variable X is the $\log_2$ of the Effector-Cell count.

(ix) *Figure S3A (Nluc activity in irradiated anti-SARS T-cell Biofactory is proportional to the number of target cells).* The Nluc activity in the irradiated or non-irradiated T-cell Biofactory stimulated by the target (*SARS-CoV-2-Sgp-cell*) or non-target (non-engineered) cells was fitted using a four-parameter logistic model, where X is the $\log_2$ of the Target-Cell count.

(x) *Figure S3B (Growth kinetics of the irradiated anti-SARS T-cell Biofactory).* A piecewise linear graph shows the T-cell Biofactory counts per mL over time (in days).

(xi) *Figures S4A – S4D (IFN production from anti-SARS T-cell Biofactory activated by SARS-CoV-1 pseudo-infected Target-Cells).* The IFN production in the T-cell Biofactory stimulated by SARS-CoV-1 pseudo-infected or non-engineered target cells was fitted using the using a four-parameter logistic model, where the independent variable X is the $\log_2$ of the Effector-Cell count.

(xii) *Figures S5A and S5B (Prophylactic activity of IFNs produced by the anti-SARS T-cell Biofactory).* The ATP activity was normalized and fitted using a four-parameter logistic model, where the independent variable X is the $\log_2$ of the IFN-dilution.

(xiii) *Figures S5C and S5D (Therapeutic activity of IFNs produced by the anti-SARS T-cell Biofactory).* The Luc2 activity, when the T-cell Biofactory was co-cultured with infected or non-infected Vero-E6-Luc2+ target cells, was fitted using a four-parameter logistic model, where the independent variable X is the $\log_2$ of the Effector-Cell count.

(xiv) *Figure S6A (Box plot showing type-I IFN signaling pathway scores).* Box plot for each treatment showing overall type-I IFN signaling pathway scores on the y-axis, using the first principal component (PC) of their expression data[56].

(xv) *Figure S6B (Heatmap of the type-I IFN signaling genes).* The normalized data was plotted by z-score. Each row of the heatmap is a single probe, and each column is a single sample (3 replicates for each treatment). Colored horizontal bars along the top of the plot identify quality control (QC) flag status and covariate categorization. Probes whose counts were below threshold in all samples were eliminated from this analysis.

(xvi) *Figures S6C and S6D (Volcano plots for differential expression of type-I IFN signaling genes).* The differential expression volcano plots were drawn using gene set analysis (GSA) and display each gene's -log10 (p-value) and log2 fold change with respect to the selected treatments. Type-I interferon signaling genes with fold change of 1.5 and adjusted p-value < 0.1 are considered significantly regulated.

(xvii) *Figures S7A – S7H (IFN signaling in Calu-3 cells treated with type-I IFNs from activated anti-SARS T-cell Biofactory).* Differential gene expression was calculated using the ΔΔCT method by normalizing the sample Ct values to mean Ct values of 3 reference genes, and expressions are represented as fold changes over untreated cell samples. Error bars represent standard errors of the mean (SEMs) from the three biological replicates.

### SUPPLEMENTARY FIGURES

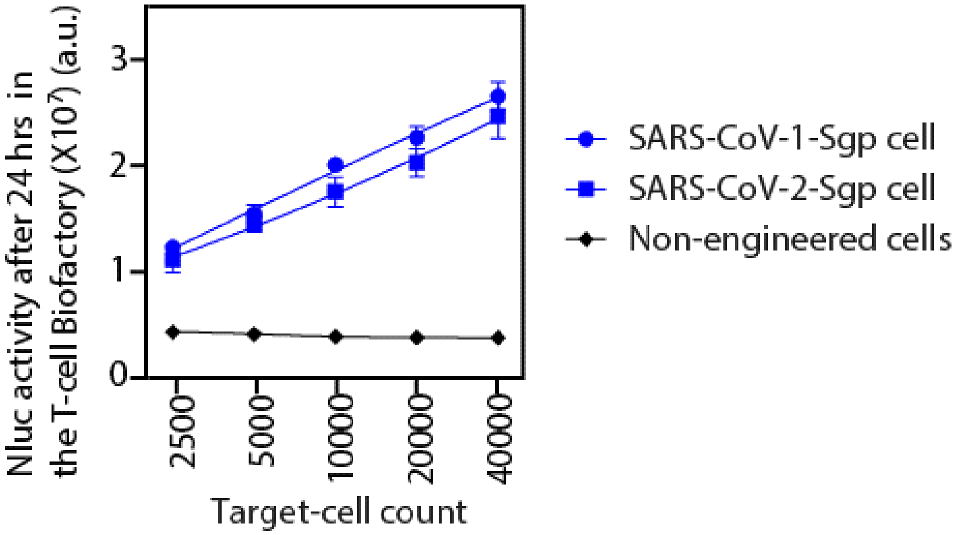


**Figure S1. Targeting SARS-CoV-2 and SARS-CoV-1 with anti-SARS T-cell Biofactory**. The Effector (Nluc) protein expression from the T-cell Biofactory is proportional to the Target-Cell count (SARS-CoV-2-Sgp-cells or SARS-CoV-1-Sgp-cells). 12,500 T-cell Biofactory were incubated with Target-Cells for 24 hours. For all observations, n = 4, error bars indicate ±1 SD.

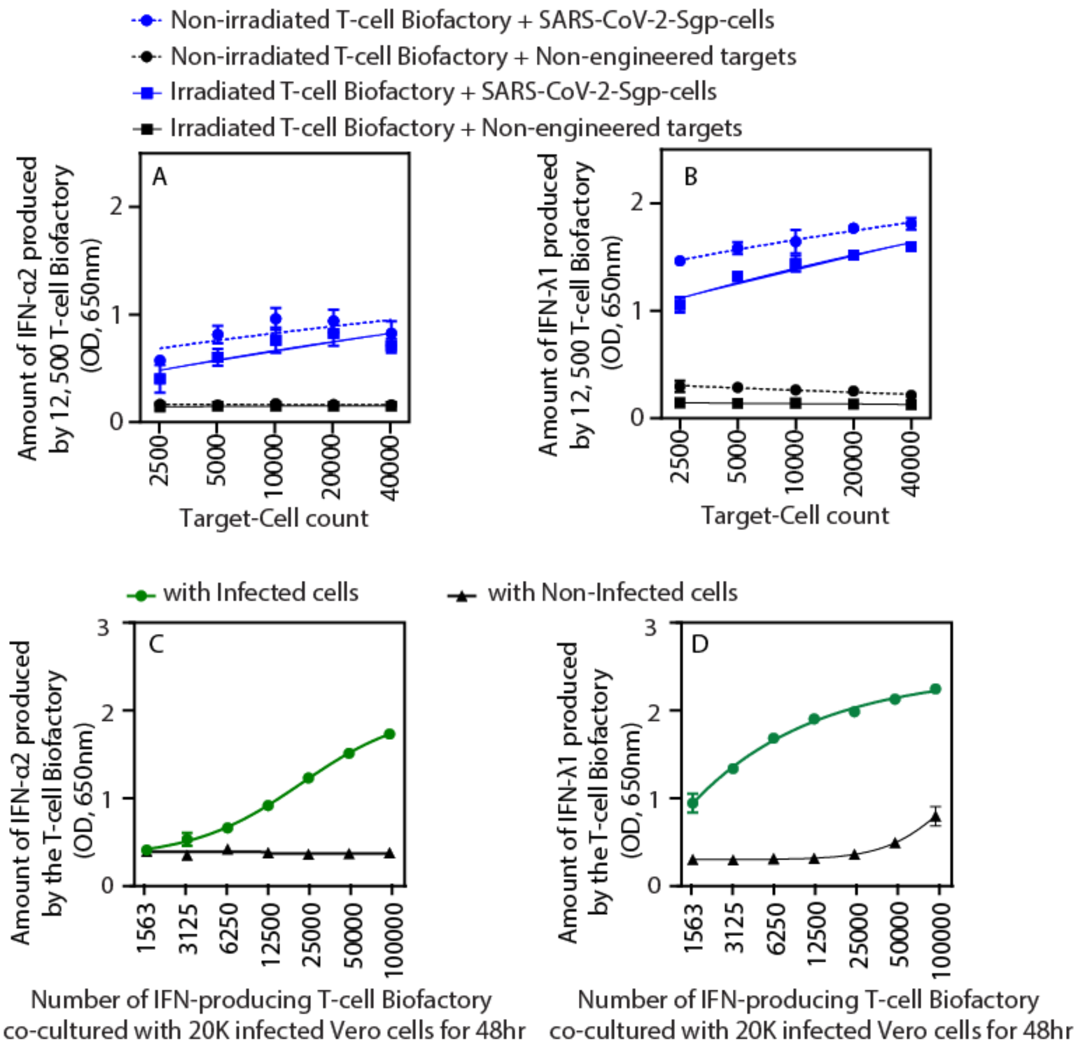


**Figure S2. IFN production by the anti-SARS T-cell Biofactory (IFN-α2, IFN-λ1).** Expression profile of type-I and type-III IFN produced from the T-cell Biofactory upon stimulation by the Target-Cells. Production of **(A)** IFN-α2 (type-I IFN) and **(B)** IFN-λ1 (type-III IFN) from the T-cell Biofactory (irradiated and non-irradiated) is proportional to the Target-Cell (SARS-CoV-2-Sgp-cells) count. 12,500 T-cell Biofactory were incubated with Target-Cells for 24 hours. Production of **(C)** IFN-α2 (type-I IFN) and **(D)** IFN-λ1 (type-III IFN) from the T-cell Biofactory is proportional to the number of T-cell Biofactories when stimulated by SARS-CoV-2-infected Vero-E6 Target-Cells. 20,000 Vero-E6 cells, infected with SARS-CoV-2 at MOI of 0.5, were used to stimulate the T-cell Biofactory for 48 hours. For all observations, n = 3, and error bars indicate ±1 SD.

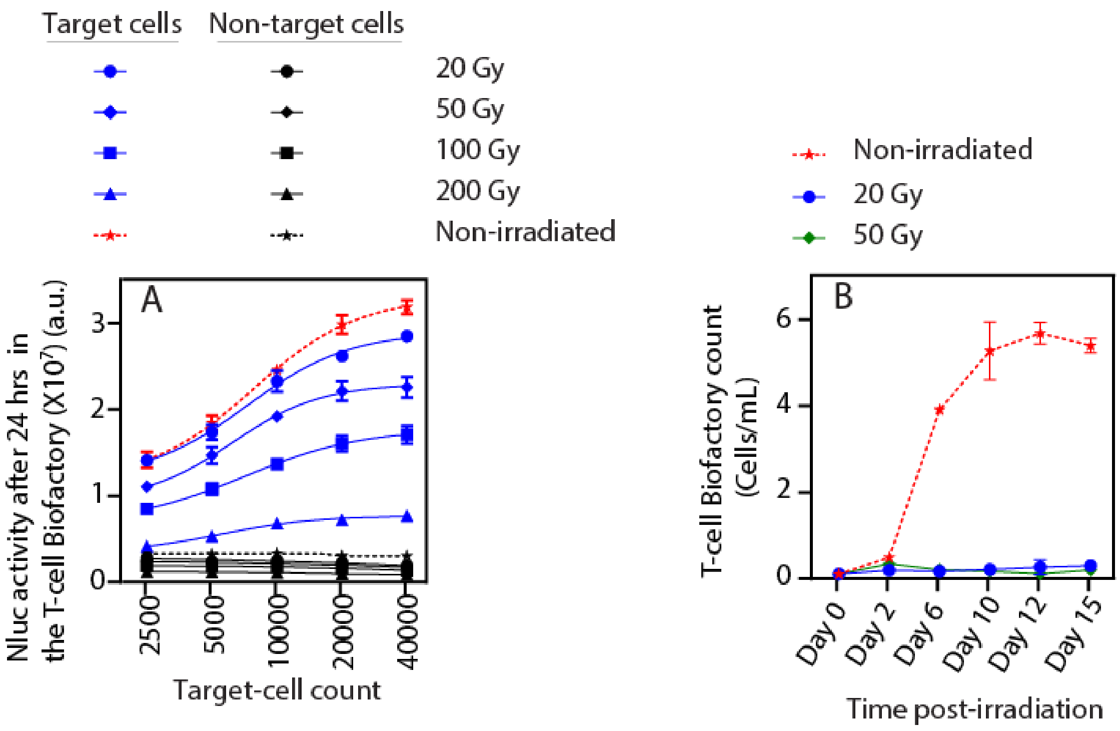


**Figure S3. Function and growth kinetics of the irradiated anti-SARS T-cell Biofactory. (A)** Effector (Nluc) protein activity in the T-cell Biofactory irradiated at different doses is proportional to the Target-Cell (SARS-CoV-2-Sgp-cells) count but reduces with the increase in radiation dose. Negative controls include the T-cell Biofactory stimulated with non-target cells, i.e., parental HEK293T/17 that do not express Sgp (non-irradiated/irradiated 12,500 T-cell Biofactory were incubated with Target-Cells for 24 hours, n = 4, and error bars indicate ±1 SD). **(B)** A dose of 20 Gy renders the T-cell Biofactory non-proliferative (n = 2, and error bars indicate ±1 SD).

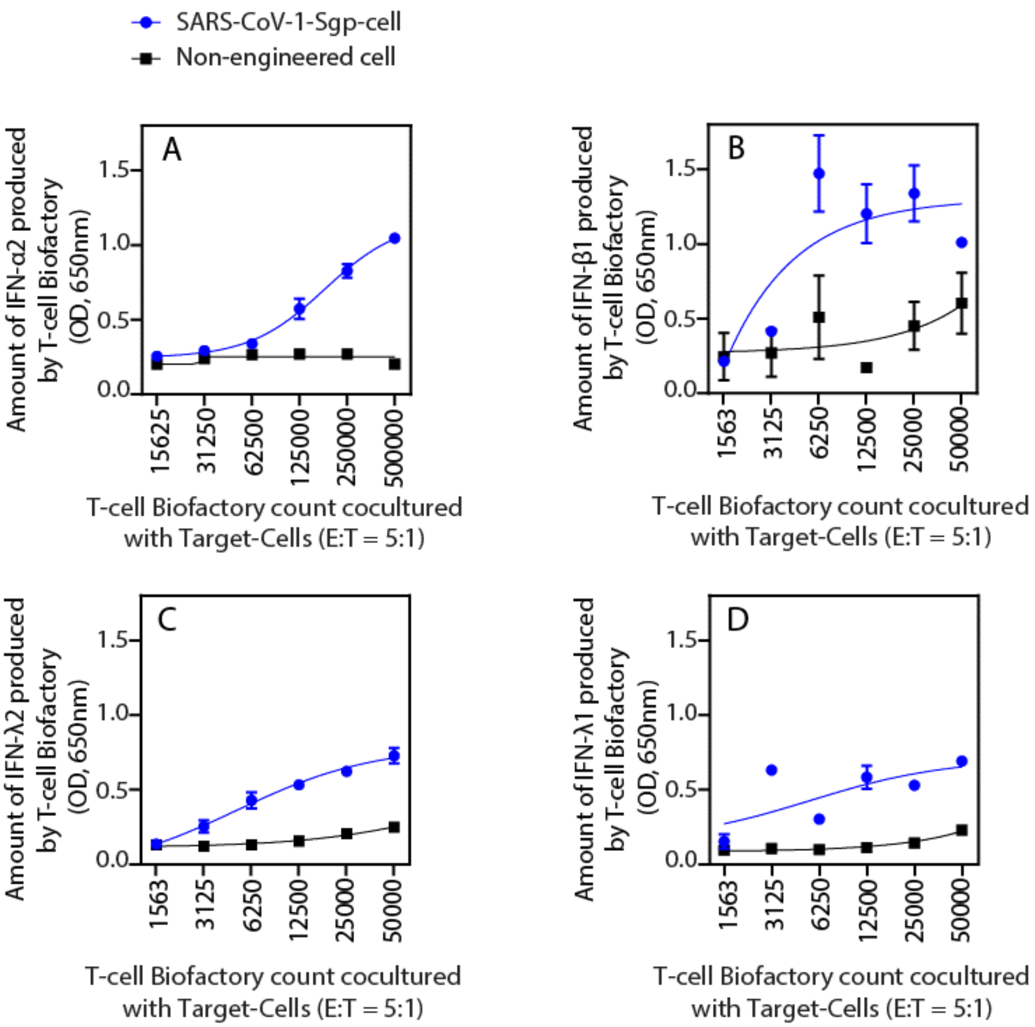


**Figure S4. Activation of the anti-SARS T-cell Biofactory by target SARS-CoV-1-Sgp-cell.** Expression profile of type-I and type-III IFN produced from the T-cell Biofactory upon stimulation by the Target-Cells. Production of type-I IFNs **(A)** IFN-α2, **(B)** IFN-β1; and type-III IFNs **(C)** IFN-λ2, and **(D)** IFN-λ1 from the T-cell Biofactory occurs upon engaging the Target-Cells (SARS-CoV-1-Sgp-cells) (E:T at all data points = 5:1, 24 hours). For all observations, n = 3, and error bars indicate ±1 SD.

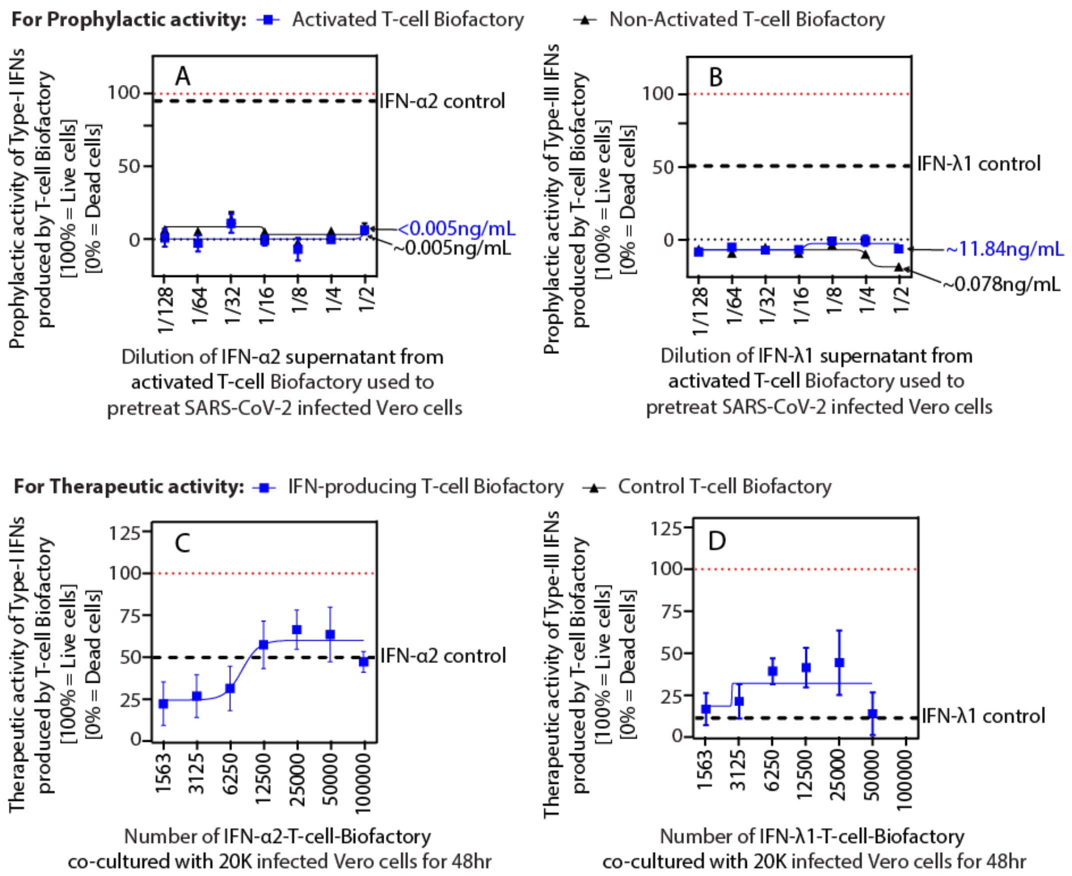


**Figure S5. Prophylactic and therapeutic activity of the anti-SARS T-cell Biofactory (IFN-α2, IFN-λ1).** The IFN produced by the T-cell Biofactory can be used as a prophylaxis **(A, B)** and a therapeutic **(C, D)** against SARS-CoV-2 infection. The protective function of the T-cell Biofactory is proportional to the amount of IFN produced **(A, C)** IFN-α2 (type-I IFN) and **(B, D)** IFN-λ1 (type-III IFN). For **(A, B)** MOI=0.1; **(C, D)** MOI=0.5. All data were collected with 20,000 Vero-E6 cells incubated with T-cell Biofactories for 48 hours. Respective recombinant human IFN at 1 µg/mL were used as controls. For all observations, n = 3, error bars indicate ±1 SD.

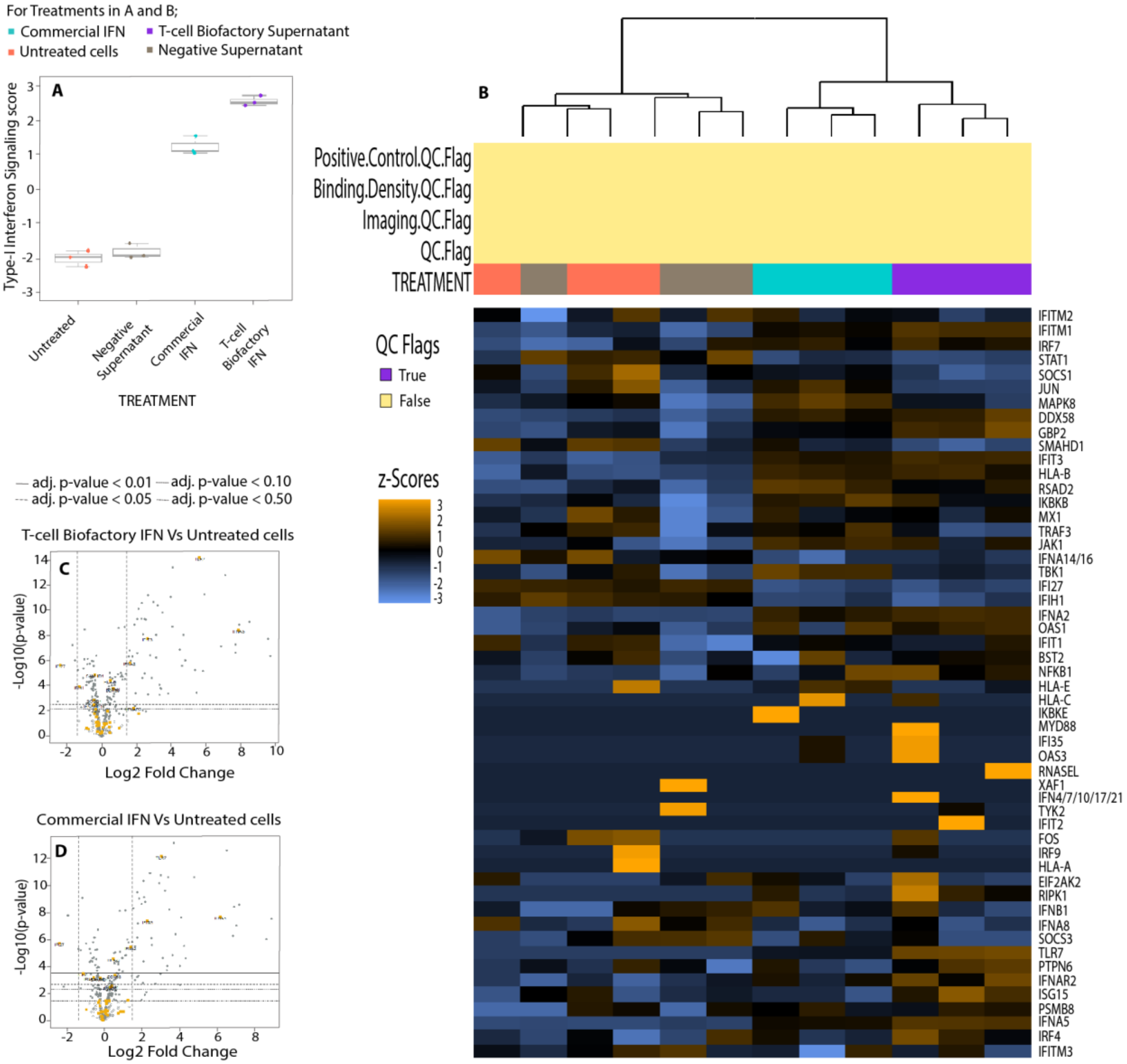


**Figure S6. Differential expression of type-I IFN signaling genes in Vero-E6 host cells using NanoString analysis.** Vero-E6 cells were treated with IFN-β1 (type-I IFN) produced from the T-cell Biofactory and assessed for regulation of the type-I IFN signaling genes using the NanoString nSolver software. **(A)** Box plot shows overall pathway scores of the type-I IFN signaling in each treatment. **(B)** Heatmap of normalized data generated via unsupervised clustering; the heatmap is scaled to give all genes equal variance. **(C, D)** Volcano plots display similar differentially expressed type-I IFN signaling gene profiles [$-\log_{10}$ (*p*-value) vs. $\log_2$ fold change]. Genes that exhibited a significant 1.5-fold change (adjusted p<0.1; vertical dotted lines) in expression in cells treated with **(C)** IFN-β1 from T-cell Biofactory and **(D)** commercial IFN-β1, compared to the control (untreated) group are shown. For all observations, n = 3, and error bars in (A) indicate ±1 SD.

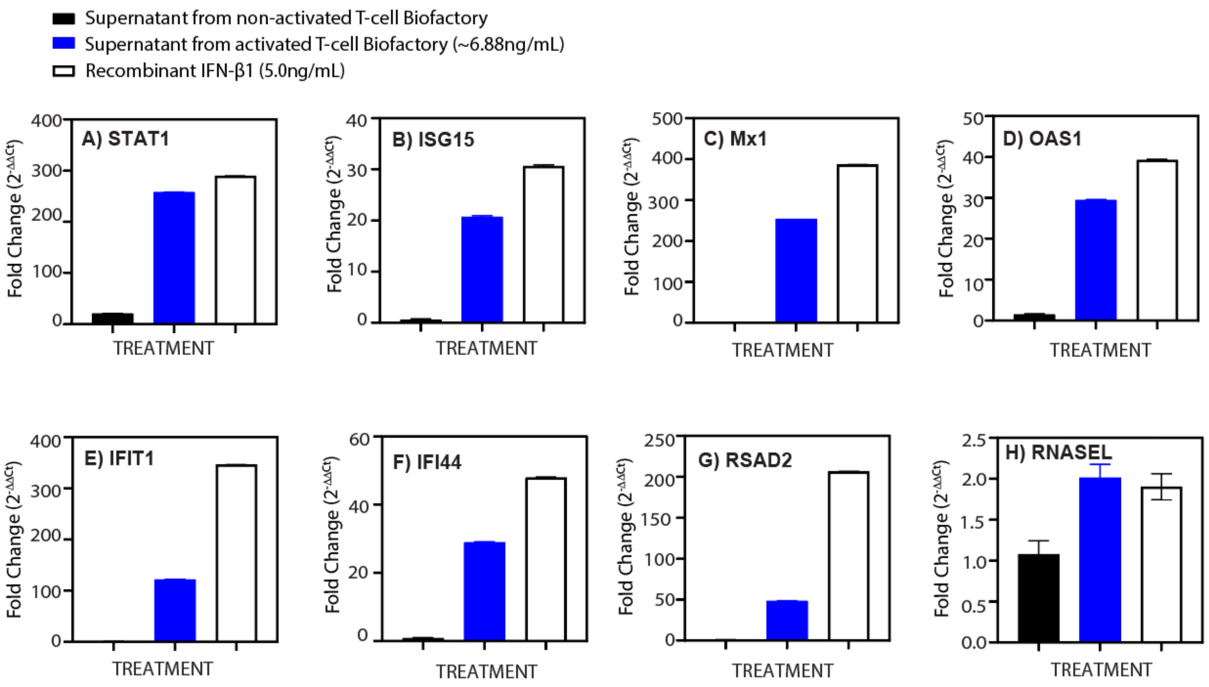


**Figure S7. Effect of IFN produced from the anti-SARS T-cell Biofactory on Calu-3 host cells.** IFN-β1 (type-I IFN) produced by the T-cell Biofactory upregulates the IFN signaling pathway, and its effect is similar to the exogenously delivered recombinant human IFN-β1 (control). Calu-3 lung epithelial cells were treated with the supernatant produced from the IFN-β1-producing T-cell Biofactory or control for 24 hours. For all observations, n = 3, error bars indicate ±1 standard errors of the mean (SEM).
